# Temperature acclimation of photosynthesis and carbohydrate metabolism are related to the geographical origin of *Arabidopsis thaliana*

**DOI:** 10.1101/2023.07.20.549837

**Authors:** Jakob Sebastian Hernandez, Dejan Dziubek, Laura Schröder, Charlotte Seydel, Anastasia Kitashova, Vladimir Brodsky, Thomas Nägele

## Abstract

Acclimation is a multigenic trait by which plants adjust photosynthesis and metabolism to cope with a changing environment. Here, natural variation of photosynthetic and metabolic acclimation was analyzed in response to low and elevated temperature. For this, 18 natural accessions of *Arabidopsis thaliana,* originating from Africa and Europe, were grown at 22°C before being exposed to 4°C and 34°C for cold and heat acclimation, respectively. Amounts of carbohydrates were quantified together with their subcellular distribution across plastids, cytosol and vacuole. Linear electron transport rates (ETRs) were determined together with maximum quantum efficiency of photosystem II (Fv/Fm) for all growth conditions and under temperature fluctuation. Under elevated temperature, residuals of ETR under increasing photosynthetic photon flux densities were found to significantly correlate with the longitudinal gradient of the geographic origin of accessions indicating a naturally occurring east-west gradient of photosynthetic acclimation capacities. Further, in heat acclimated plants, vacuolar fructose amount was found to positively correlate with longitude while plastidial and cytosolic amounts were found to be negatively correlated. Plastidial sucrose concentrations were found to positively correlate with maximal ETRs under fluctuating temperature indicating a stabilizing role within the chloroplast. In summary, our findings revealed specific subcellular carbohydrate distributions which contribute differentially to photosynthetic efficiencies of natural *Arabidopsis thaliana* accessions across a longitudinal gradient. This sheds light on the relevance of subcellular metabolic regulation for photosynthetic performance in a fluctuating environment and supports the physiological interpretation of naturally occurring genetic variation of temperature tolerance and acclimation.

## 1. Introduction

Environmental dynamics have direct effects on plant metabolism and performance. Plant stress response and acclimation to environmental changes stabilize the physiological homeostasis and prevent irreversible tissue damage or adverse effects on development and growth. Both stress response and acclimation are multigenic traits and typically comprise many physiological and molecular changes, including significant reprogramming of photosynthesis, primary and secondary metabolism (Hannah et al., 2006; Garcia-Molina et al., 2020; Schwenkert et al., 2022; Seydel et al., 2022). For example, a changing temperature regime affects properties of membrane systems and enzyme kinetics resulting in shifts of membrane fluidity, transmembrane concentration gradients and reaction rates (Elias et al., 2014; Cano-Ramirez et al., 2021). Metabolic pathways and signaling cascades need to be reprogrammed in order to sustain and stabilize metabolism (Herrmann et al., 2019). If environmentally driven deflection from a metabolic state exceeds a certain threshold, this might lead to the formation of reactive oxygen species (ROS) and irreversible tissue damage (Choudhury et al., 2017). Below this threshold, plants are able to prevent irreversible damage by adjusting metabolism, membrane structure and the composition of the photosynthetic apparatus. For example, low temperature results in an increase of cytochrome b6f complex, ATP synthase, Rubisco and other Calvin-Benson-Bassham cycle (CBBC) enzymes whereas heat leads to increased proportions of light harvesting complex II (LHCII) and photosystem I (PSI) while CBBC activity is downregulated (Gjindali and Johnson, 2023). This shows that changes of environmental temperature regimes cause diverse adjustments of various processes on a molecular and physiological scale which challenges our understanding and experimental analysis of pathways which are involved in temperature stress response and acclimation. Further complication of experimental and theoretical analysis is added on by interaction effects of different environmental stimuli, e.g., effects different light intensities have on temperature response (Huner et al., 1998). Although light has been found to be essential for full acclimation to low temperature (Wanner and Junttila, 1999), too high intensities result in photoinhibition and photooxidative damage (Gray et al., 2003).

When exposed to low temperature, sucrose metabolism was observed to play a key role in stabilizing photosynthetic efficiency, CO_2_ fixation and carbon allocation (Kitashova et al., 2023). Before it was shown that carbon uptake in a low temperature regime is limited by the capacity to synthesize sucrose (Strand et al., 2003). Cold sensitive natural accessions of *Arabidopsis thaliana* were found be restricted in their maximal sucrose phosphate synthase (SPS) activity which might result in ROS generation (Nägele et al., 2012). Furthermore, photosynthesis was found to be stabilized by vacuolar sucrose cleavage as metabolic control is preserved under cold stress (Weiszmann et al., 2018; Nägele, 2022). These findings suggest a central role of subcellular sucrose metabolism for temperature stress response and acclimation. Previously, it has been suggested that carbon partitioning plays a crucial role in enhancing photosynthesis at low temperatures. Specifically, a higher flux of carbon directed towards sucrose, rather than starch, can result in this improvement (Lundmark et al., 2006). A theoretical model has demonstrated that a shift in carbon partitioning towards sucrose can stabilize metabolism in the face of fluctuating environmental conditions (Hernandez and Nägele, 2022). In addition to changing carbon partitioning, increased starch degradation results in a drop of starch amount during the first 24 h of cold exposure. This was suggested to serve as carbon resource for hexose biosynthesis which have the capacity to act as cryoprotectants and a source for rapid energy supply (Sicher, 2011). Following this initial decrease, plants then accumulate starch to higher levels as before cold exposure (Guy et al., 2008). It was reported earlier that natural variation of starch accumulation during cold acclimation might be explained by a differential regulation of the starch degradation pathway resulting in higher starch amount in the freezing sensitive accession Cvi-0 (Cape Verde Islands), originating from Africa, than in the freezing tolerant accession Rsch (Rschew), originating from Russia (Nagler et al., 2015). Comparing Rsch under elevated temperature to the Sicilian accession Ct-1 (Catania) revealed that, similar to cold, Rsch accumulated lower starch amount than Ct-1 (Atanasov et al., 2020). However, in contrast to cold, total starch amount decreased under heat compared to control conditions, and the decrease was stronger in Rsch than in Ct-1. Further, decreased starch amount under heat is rather due to inhibition of ADP-glucose pyrophosphorylase and starch synthesis than increased capacities of starch degradation (Geigenberger et al., 1998; Awasthi et al., 2014).

Recently, availability of sugars has been shown to be essential for plant survival under heat (Olas et al., 2021). While this demonstrates the crucial role of carbohydrate metabolism, its regulation under such conditions is less well understood. It has become evident that, to comprehensively address questions about metabolic regulation, subcellular and compartment-specific data is required (Nägele and Heyer, 2013; Hoermiller et al., 2017; Fürtauer et al., 2019; Höhner et al., 2021). For example, it was found that at low temperatures substantial reorganization of soluble sugar concentrations occur (Hoermiller et al., 2017; Weiszmann et al., 2018; Patzke et al., 2019). This subcellular reorganization is hypothesized to be both necessary to distribute cryoprotective sugars to key cellular structures, e.g., raffinose to thylakoid membranes, and to stabilize the cellular metabolic homeostasis (Knaupp et al., 2011; Nägele and Heyer, 2013). Also, under elevated temperature, subcellular compartmentation and reprogramming of metabolism plays an important role, e.g., for regulation of signaling cascades and ROS production (Kohli et al., 2019). Changes in subcellular signaling might explain the previously observed downregulation of transcripts of photosynthesis and carbohydrate metabolism under heat stress (Prasch and Sonnewald, 2013). While transcripts are downregulated, sugar amounts were found to either stabilize or increase during heat acclimation (Atanasov et al., 2020; Garcia-Molina et al., 2020). Simultaneously, a heat-induced increase of neutral and cell wall-associated invertase activities may explain an observed hexose accumulation which was more pronounced in a natural *Arabidopsis* accession originating from southern Europe than in a northern accession (Atanasov et al., 2020).

In summary, these findings indicate that it is necessary to study carbohydrate metabolism on a subcellular level which is hardly predictable from the genome or transcriptome. With such an approach, regulatory patterns might be identified which stabilize photosynthesis in a changing temperature regime. Comparing natural accessions of *Arabidopsis thaliana* has been shown before to be a promising strategy to unravel conserved and specialized stress response and acclimation mechanisms (Maloof et al., 2001; Hannah et al., 2006; Weiszmann et al., 2023). Here, we analyzed the role of subcellular sugar compartmentation for stabilization of photosynthetic efficiency by comparing natural variation of metabolic acclimation in *Arabidopsis* to both a high and a low temperature regime. Natural *Arabidopsis* accessions, originating from a wide geographical range, were acclimated to low and high temperature before being exposed to a fluctuating temperature regime. Carbohydrates were quantified on a cellular and subcellular level. Finally, we developed an app to estimate subcellular compound distribution from experimental data using Monte Carlo simulations.

## 2. Material and Methods

### 2.1 Growth conditions and plant material

Plants of 18 natural accession of *Arabidopsis thaliana* were grown for 5 weeks in a climate chamber under short day conditions (8 h/16 h light/dark; photosynthetically active radiation (PAR): 100 µmol photons m^-2^ s^-1^; 22 °C/16 °C; 60% relative humidity; see Supplementary Table S1 for a full list of natural accessions). Then, with same daylength, PAR and relative humidity, plants were transferred to 4°C, 34°C, or left at 22°C for a duration of 4 days.

Plant material was sampled at midday, i.e., after 4 hours of light exposure. Leaf rosettes were cut with a scalpel at the hypocotyl and immediately transferred to liquid nitrogen to quench metabolism. Under constant supply of liquid nitrogen, plant material was ground to a fine powder, and subsequently lyophilized.

### 2.2 Chlorophyll fluorescence measurements

Light response curves were recorded using a pulse-amplitude-modulation (PAM) protocol on single leaves using a WALZ JUNIOR-PAM® (www.walz.com) in which photosynthetic photon flux density (PPFD, µmol photons m^-2^ s^-1^) was stepwise increased every 20 sec after 15 min of dark incubation at 4°C, 22°C or 34°C for all acclimation conditions (PPFD list: 0, 40, 72, 104, 144, 200, 304, 456, 672, 1000, 1312, 1840, 2400 µmol photons m^-2^ s^-1^). This achieved quantifying photochemical energy conversion efficiency in acclimated plants (growth temperature = measurement temperature) and under temperature stress (growth temperature ≠ measurement temperature). The linear electron transport rate (ETR) was calculated as follows (Eq. 1):

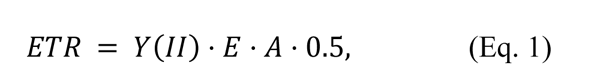

with *Y*(*II*) being the effective photochemical quantum yield of PS II, *E* the incident irradiance, and *A* the absorptance.

### 2.3 Quantification of carbohydrates

Extraction and determination of starch and soluble sugar concentrations was done as described before (Kitashova et al., 2023). In brief, lyophilized leaf material was incubated twice with 80% ethanol at 80°C for 30 min to extract soluble sugars. The pellet contained leaf starch granules which were hydrolyzed, digested with amyloglucosidase, and quantified in a photometric assay using a coupled reaction of glucose oxidase, peroxidase and o-dianisidine. Extracted supernatants were pooled, dried in a desiccator and sugars in the pellet were dissolved in H_2_O_dd_. Amounts of the soluble carbohydrates sucrose, glucose and fructose were determined photometrically. Sucrose was determined after incubation with 30% KOH at 95°C using an anthrone reagent composed of 14.6 M H_2_SO_4_ and 0.14% (w/v) anthrone yielding a complex with a specific absorbance maximum at 620 nm. Glucose and fructose amounts were determined within a coupled hexokinase/glucose 6-phosphate dehydrogenase assay which yielded NADPH + H^+^ detectable at 340 nm.

### 2.4 Non-aqueous fractionation (NAF)

Subcellular fractionation followed a NAF protocol described earlier (Fürtauer et al., 2016). After the fractionation of lyophilized leaf material in mixtures of tetrachlorethylene and n-heptane with differential densities, marker enzyme activities were determined for plastids (alkaline pyrophosphatase activity), cytosol (UDP-glucose pyrophosphorylase activity) and vacuole (acidic phosphatase activity). Subcellular amounts of sucrose, glucose and fructose were determined as described in the previous paragraph.

To calculate the relative distributions of metabolites present in each subcellular compartment, an approach based on linear regression and a Monte Carlo simulation was used. This approach is based on the fact, that metabolite distributions are made from a linear combination of the compartment-specific gradients (Eq. 2):

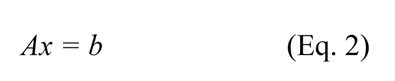

With *A* being the matrix containing relative values of the marker metabolites or enzymes. Each row *m* represents a density fraction and each column *n* a subcellular compartment. The vector *b* consists of the relative distribution of our target metabolite present in each compartment *x* and is the unknown vector of interest. Finally, the vector *b* consists of the relative values of target metabolites in each fraction *m*.

Thus, to obtain the relative distribution of a target metabolite across the subcellular compartments, (Eq. 2) needs to be solved for *x*. For this, linear least-squares fitting constrained to the interval [0, 1] was applied, as a relative distribution within a compartment cannot be negative or greater than 1.

However, experimental data has shown that a technical error of 10% cannot be excluded when performing NAF (Fürtauer et al., 2016). To account for this, random noise was added to each measurement following a normal distribution with a standard deviation of 5%. This resulted in more robust results and negates infeasible relative distributions such as *x* = (0, 0, 1)^*T*^.

This method has been converted into an R Shiny App for ease of use and can be found in a GitHub repository (https://github.com/cellbiomaths/NAFalyzer). The app also contains a detailed explanation of how to apply and validate NAF data.

### 2.5 Climate data

Climate data was obtained from a database (https://climatecharts.net/) and comprises values from 1950 until 2019. Temperature mean of a month is given in [°C] and precipitation in [mm month^-1^].

### 2.6 Statistics and data evaluation

Statistics and data evaluation was performed in R (Version 4.2.2) using RStudio, Version 2022.07.2 Build 576 (R Core Team, 2021).

## 3. Results

### 3.1 Fluctuation of linear ETR reveals a longitudinal gradient of natural accessions

After 5 weeks of growth under short day conditions, plants of 18 natural accessions were acclimated to low (4°C) or elevated (34°C) temperature. Maximum quantum yield of PSII (Fv/Fm) and linear ETR were quantified to reveal photosynthetic efficiencies. Detailed information about the accessions is provided in the supplement (Supplemental Table S1). Measurements were conducted at the temperature which plants were acclimated to, i.e., 4°C, 22°C and 34°C, and also in cross-conditions to reveal the effect of a sudden change in temperature regimes on photosynthesis (Supplemental Figure S1). Linear ETR was found to be both dependent on the measuring and the acclimation temperature. Plants acclimated and measured under 34°C showed the highest mean rates, while plants acclimated at 4°C and measured at 4°C demonstrated the lowest mean rates of ETR.

Particularly, yet not exclusively, from cross-conditional measurements, fluctuations of ETR curves were recognized, which indicate instability in photosynthesis over increasing light intensities. To quantify these fluctuations, a curve following a basic saturation kinetic was fitted to the data (Eq. 3):

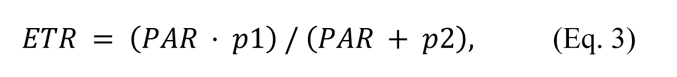

with *p1* and *p2* being optimized parameters. The resulting residuals between the fit and experimental data then represent a measure for the degree of fluctuation of ETR. Strongest fluctuations were observed at PAR > 500 µmol photons m^-2^s^-1^ while for PAR < 500 µmol photons m^-2^s^-1^ only a weak signal-to-noise ratio was identified (see Supplemental Figure S1). Due to this, residual analysis focused on ETR for PAR > 500 µmol photons m^-2^s^-1^. Further, for scaling across measurements, all residuals were normalized to the maximum ETR observed in the respective measurement.

Mean scaled residuals across all accessions became highest for plants acclimated and measured at 4°C (Fig. 1 A). In general, residuals were observed to be higher for measurements at 4°C than at 22°C and 34°C. Yet, interestingly, cold acclimation resulted in higher residuals than in non-cold or heat acclimated plants when measured at 4°C. In contrast, heat and cold acclimation resulted in a lower fluctuation level of ETR compared to non-acclimated plants (22°C) when measured at 34°C (Fig. 1 A, right panel). To test whether scaled residuals can provide information about geographic origin and habitat of natural accessions, they were Spearman correlated with latitude, longitude, temperature, and precipitation of the first quarter of the year (Fig. 1 B). This part of the year has recently been found to be indicative of regulation of primary metabolism of natural *Arabidopsis* accessions (Weiszmann et al., 2023).

**Fig. 1.**
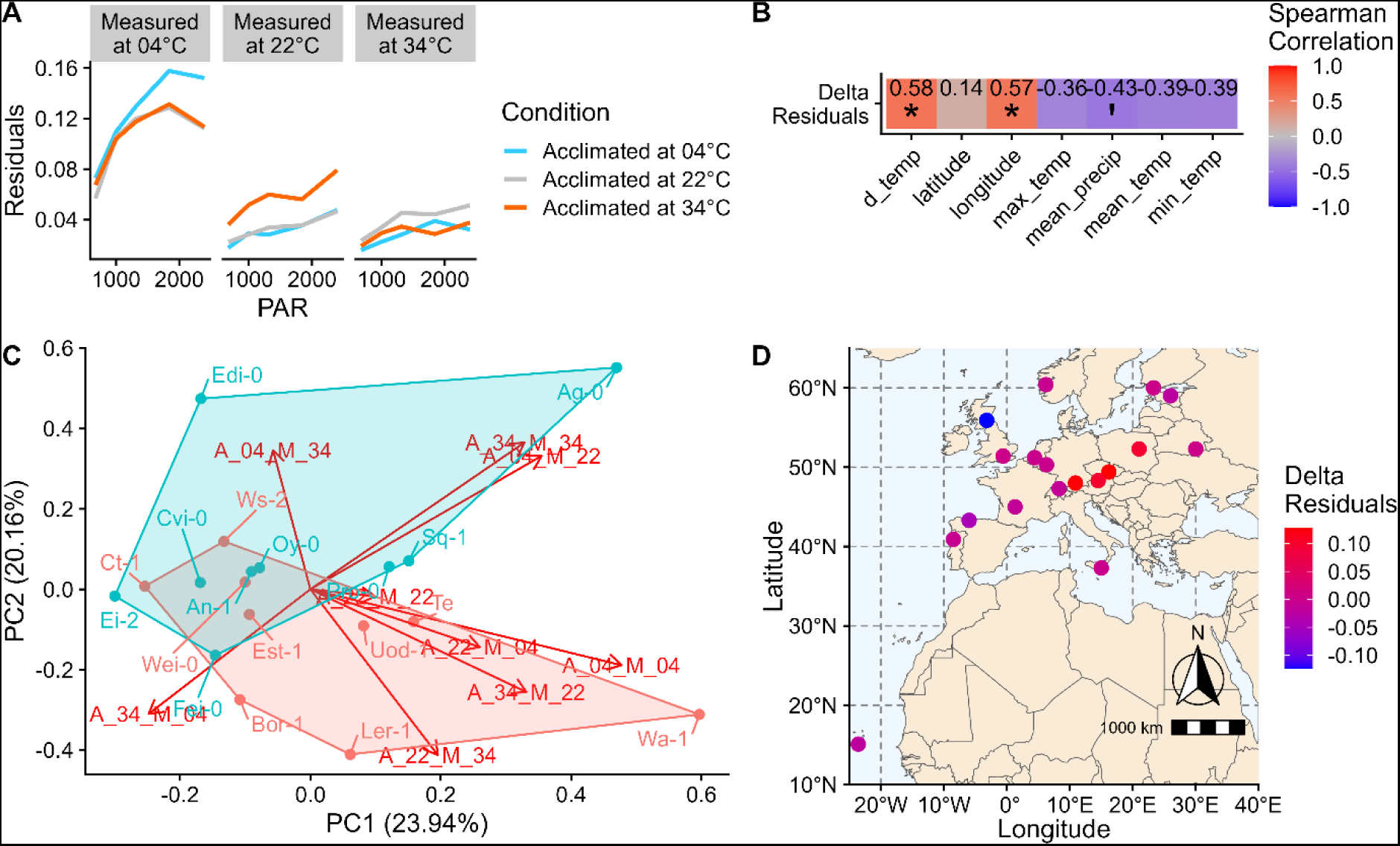
Fluctuations of electron transport rates. **(A)** Overall trends of residuals in each combination of growth and measurement temperatures across all natural accessions. Residuals were normalized to the maximum ETR. (**B)** Spearman correlation between geographical and climate data to the difference between the mean scaled residuals of plants measured at 34°C which were acclimated at 22°C and 04°C. Climate data comprise the first quarter of year (i.e. Jan – Mar). d_temp: maximum temperature difference; max_temp: highest temperature; mean_precip: mean precipitation; mean_temp: mean temperature; min_temp: minimal temperature. Details are provided in the supplements (Supplemental Table S1). Asterisks indicate significance: * p < 0.05. Mean precipitation was significantly correlated with residuals to a significance level of 10 %, i.e., p < 0.1 (**‘**). Red colour indicates positive correlation, purple colour indicates negative correlation. (**C)** PCA of the mean scaled residuals of all conditions. Accessions labeled in blue represent the accessions classified as ‘western’, while the red label represents ‘eastern’ accessions. Loadings indicate mean scaled residuals at a certain acclimation (A) and measurement (M) temperature, e.g.: *A_04_M_34: acclimated at 4°C, measured at 34°C*. (**D)** Geographic origins of accessions and the delta in residuals of plants acclimated at 22°C and 04°C, when measured at 34°C.

When measured at 22°C, plants acclimated to 4°C and 22°C showed similar residuals of ETR, while plants acclimated at 34°C showed the highest fluctuations. When measured at 34°C, plants acclimated at 4°C and 34°C showed most stable ETR while 22°C showed higher scaled residuals. This observation indicated a conserved mechanism of temperature acclimation and stabilization of photosynthetic efficiency. Hence, for further temperature acclimation analysis, the difference between the mean scaled residuals of plants acclimated at 22°C and 4°C, both measured at 34°C, was calculated and described as *Δ(delta) residuals* indicating the acclimation capacity of photosynthetic electron transport under maximal temperature fluctuation. The *Δresiduals* were correlated to geographical and climate data from the first quarter of the year at the original habitat of the accessions. The analysis revealed a significant correlation between the *Δresiduals* and the longitude, as well as the difference in monthly mean temperatures recorded within the first quarter of the year, i.e., *d_temp = maximum mean temperature – minimum mean temperature*. Due to the significant positive correlation of *Δresiduals* with longitude, accessions were classified as western (50%, i.e., 9 out of 18 accessions) or eastern (50%; Fig. 1 D). Principal component analysis (PCA) showed that variance on PC1 was explained by measurements at 4°C while PC2 was explained by measurements at 34°C (Fig. 1 C). Most distant accessions on PC1 were Ei-2 (Eifel, Germany) and Wa-1 (Warsaw, Poland). Most distant accessions on PC2 were Ag-0 (Argentat, France) and Ler-1 (Landsberg am Lech, Germany).

Maximum quantum yield of PSII (Fv/Fm) did not significantly differ between eastern and western accessions under neither analyzed condition (Supplemental Figure S1). In general, it was lowest for plants acclimated to 4°C (Fv/Fm ∼0.7-0.8) and highest for non-acclimated plants grown and analyzed at 22°C (Fv/Fm ≥0.8). Variance of Fv/Fm was highest for cold acclimated plants suggesting higher variability of acclimation of PSII among natural accessions than under elevated temperature. The Fv/Fm showed significant negative correlation with ETR residuals, except for non-acclimated plants or heat acclimated plants when measured at 4°C (Supplemental Figure S2). Finally, short-term cold treatment induced higher photosynthetic fluctuations than both long-term and short-term heat exposure.

### 3.2 Differential carbohydrate accumulation during heat and cold acclimation relates to a longitudinal gradient

Amounts of starch, sucrose, glucose, and fructose were quantified across all 18 natural accessions under each acclimation condition. Soluble carbohydrates significantly increased during cold acclimation, but total amounts varied strongly between accessions (Supplemental Figure S3). In heat acclimated plants, sucrose amount also increased but to a lower extent than in the cold whereas hexose amounts decreased compared to non-acclimated plants. The sucrose to starch ratios in C6 equivalents showed distinct patterns across the natural accessions under low and high temperature. For most accessions, the ratio of sucrose to starch was higher when plants were exposed to heat rather than cold. In Pro-0, Wei-0, and Cvi-0, the sucrose to starch ratio in cold treated plants remained similar to non-acclimated plants.

In a PCA, temperature-induced carbohydrate dynamics separated cold acclimated plants from non-acclimated (22°C) and heat acclimated plants on PC1 explaining almost 80 % of total variance (Figure 2). On PC2, a separation of eastern and western accessions was observed which was strongest for cold acclimated plants and weakest for non-acclimated plants. A differential ratio of starch to soluble carbohydrates explained the observed separation on PC2 (14%). In general, starch accumulation was stronger in western accessions while soluble carbohydrates were accumulated stronger in eastern accessions.

**Fig. 2.**
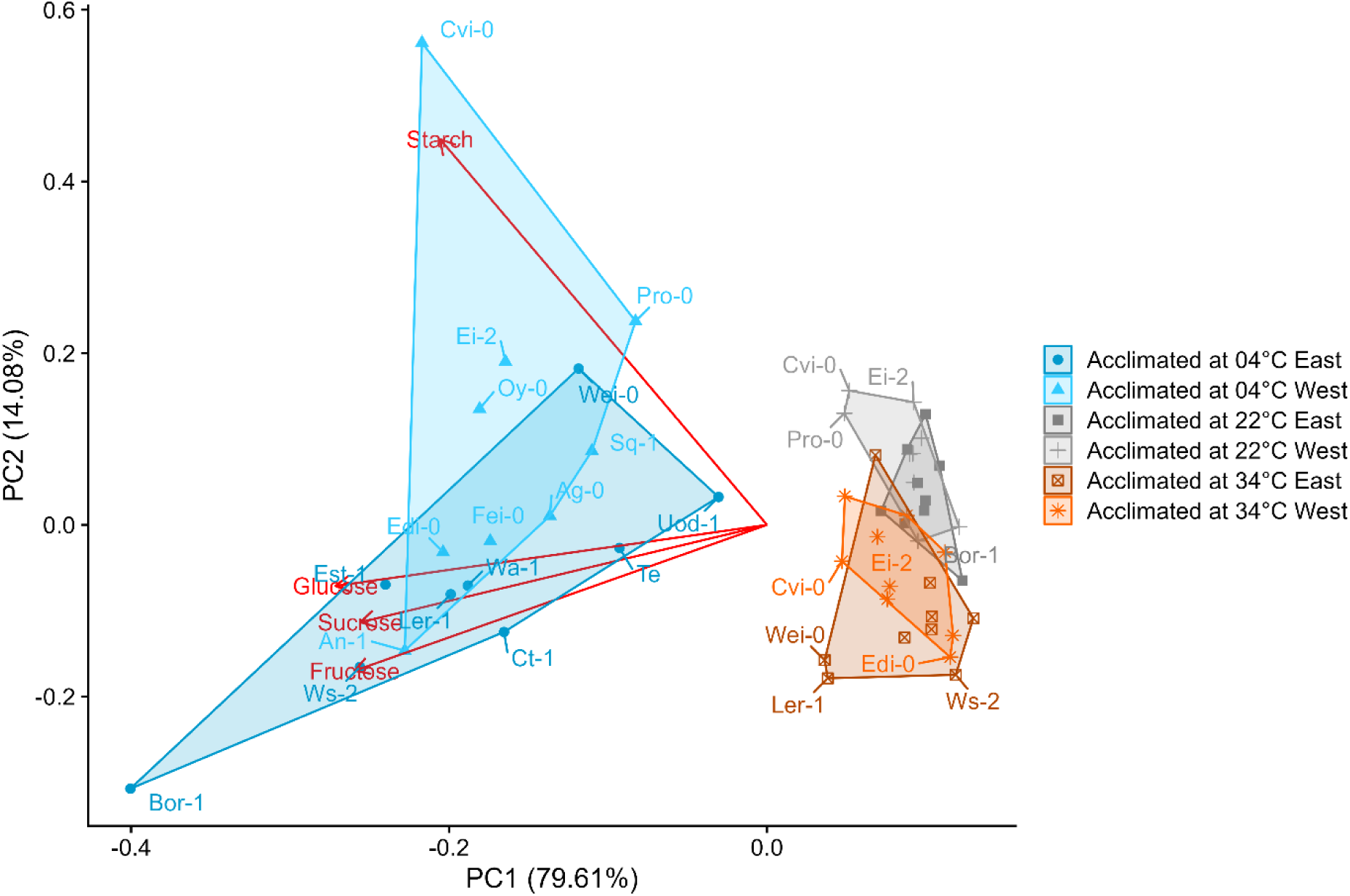
Differential carbohydrate accumulation during heat and cold acclimation in eastern and western accessions. Dots represent scores of accessions acclimated at different temperatures as indicated by the color (grey: 22°C, blue: 4°C, orange: 34°C). Loadings, i.e., carbohydrates, are shown in red (n ≥ 5). Classification of eastern and western accessions refers to scaled ETR residuals of heat acclimated plants (see Figure 1).

While for starch the subcellular localization is defined as the chloroplast, metabolism of soluble sugars is hardly predictable by their total amount. Subcellular distribution of sucrose, glucose and fructose was determined experimentally applying a non-aqueous fractionation protocol which enables the separation of plastidial, cytosolic and vacuolar fractions while metabolism is continuously quenched (Fürtauer et al., 2016). A conserved cold response across accessions was an increased proportion of sucrose in plastids and cytosol while the vacuolar proportion decreased compared to plants at 22°C (Fig. 3 A). At 34°C, only cytosolic portions increased while vacuolar sucrose content decreased. The plastidial relative sucrose content remained similar to 22°C plants. Relative glucose fractions were highest in the vacuole across all conditions and accessions, and both low and elevated temperature resulted in an even stronger vacuolar accumulation (Fig. 3 B). For fructose, a similar distribution was observed except for plants acclimated to 34°C (Fig. 3 C). Here, it was observed that the median fructose portion increased in the plastids and decreased in the vacuole. Further, fructose compartmentation at 34°C was most variable across accessions ranging from a similar distribution across all compartments in Cvi-0 to a very strong vacuolar compartmentation in Ct-1, Est-1 and Fei-0 (Fig. 3 C, lower panel).

**Fig. 3.**
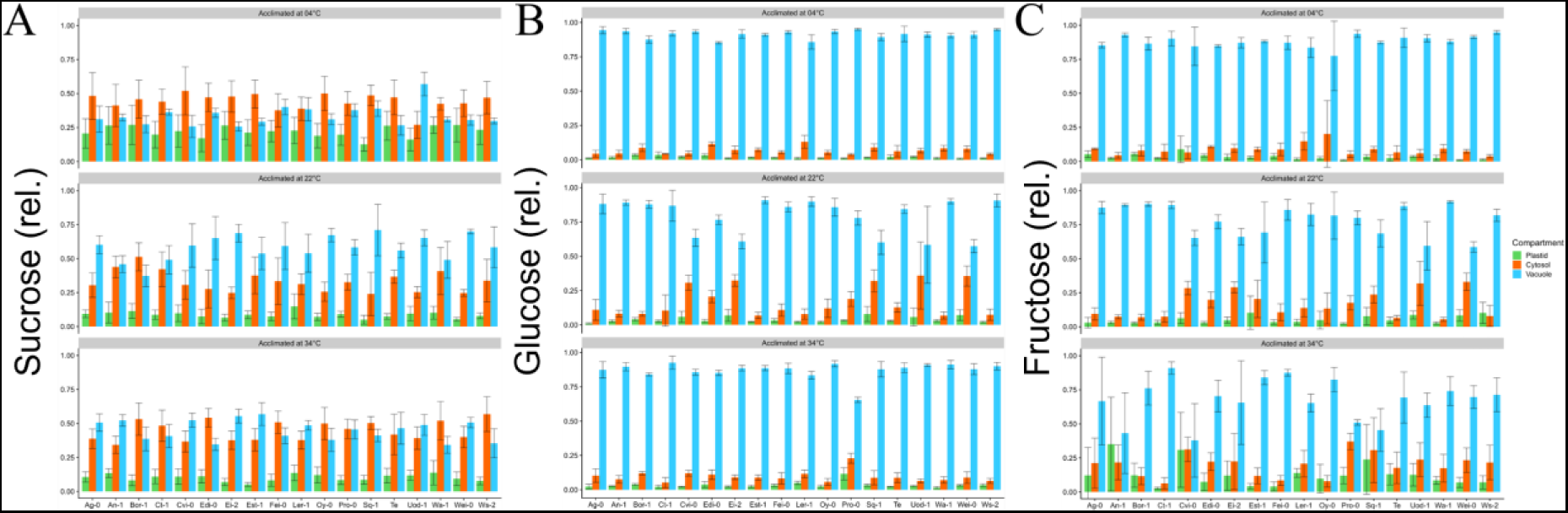
Subcellular carbohydrate compartmentation of cold and heat acclimated natural accessions. (**A**) Relative sucrose amounts of plastids (green), cytosol (orange) and vacuole (blue). Top panel: plants acclimated at 4°C. Middle panel: plants acclimated at 22°C. Lower panel: plants acclimated at 34°C. (**B**) Relative glucose amounts of plastids (green), cytosol (orange) and vacuole (blue). Top panel: plants acclimated at 4°C. Middle panel: plants acclimated at 22°C. Lower panel: plants acclimated at 34°C. (**C**) Relative fructose amounts of plastids (green), cytosol (orange) and vacuole (blue). Top panel: plants acclimated at 4°C. Middle panel: plants acclimated at 22°C. Lower panel: plants acclimated at 34°C. Ordinate: relative metabolite amount (0-1); abscissa: accessions. Bars represent means ± SD (n = 3).

To test whether fructose compartmentation was related to the geographical origin or habitat information of accessions, subcellular amounts were correlated with temperature, longitude and latitude (Table I). Cold acclimated plants showed a significant positive correlation of plastidial fructose with the mean temperature of the first quarter of the year at the place of geographic origin. Longitude and latitude both were correlated negatively with the relative plastidial fructose amount (Pearson, 0.001 < p < 0.05). For non-acclimated plants (22°C) no significant correlation was observed to a significance level of 10% (Pearson, p > 0.1). For heat acclimated plants, vacuolar fructose amount was found to be positively correlated to longitude (Pearson, 0.01 < p < 0.05) while plastidial and cytosolic amounts were negatively correlated (Pearson, 0.05 < p < 0.1).

**Table I:**
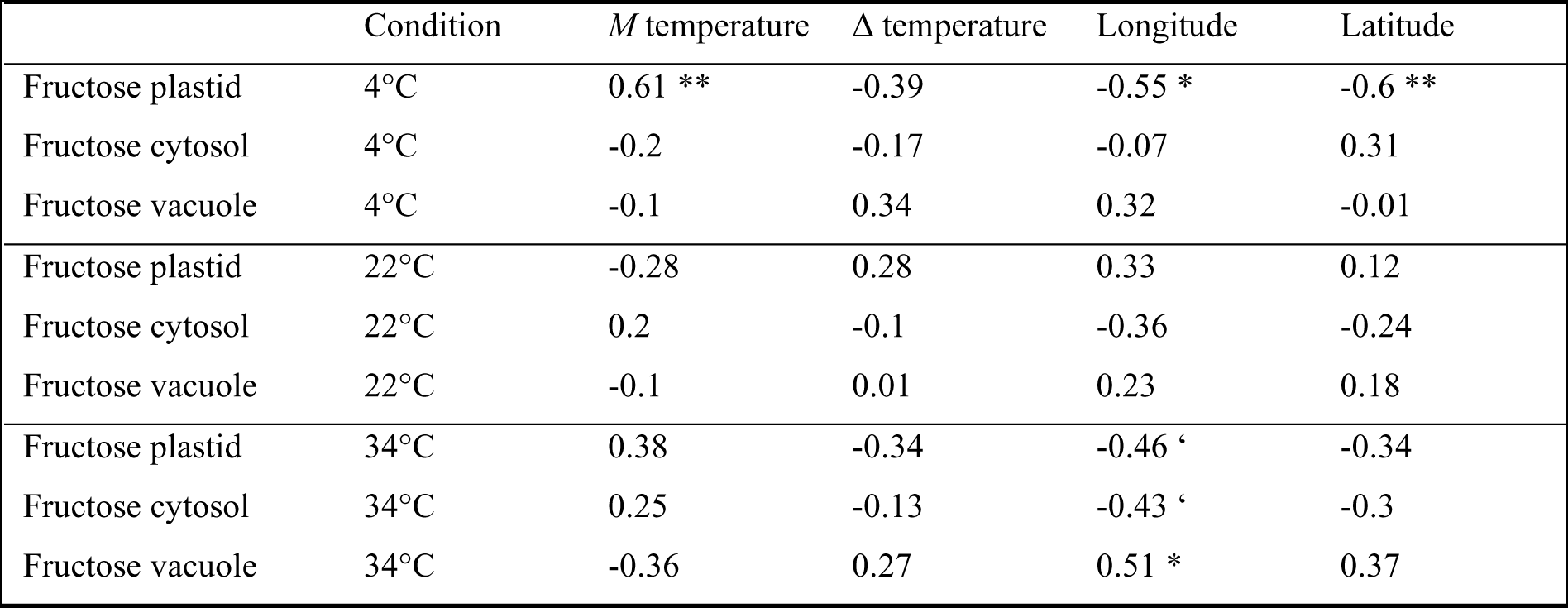
Pearson correlation coefficients of subcellular fructose distribution, climate and geographical data of natural accessions. Climate data was used from the first quarter of the year, i.e., January to March. *M temperature* mean temperature; *Δ temperature:* difference between the highest and lowest average temperature of the individual months. Significance codes: ** 0.001 < p < 0.01; * 0.01 < p < 0.05; ‘ 0.05 < p < 0.1. Information about glucose and sucrose is provided in the supplements (Supplemental Table S2).

While relative fructose amount in the cytosol and plastids increased in heat acclimated plants, vacuolar hexose compartmentation and plastidial sucrose allocation was found to separate cold acclimated accessions from heat and non-acclimated plants (Fig. 4). Plastidial sucrose allocation was found to be a conserved cold response across western and eastern accessions. In contrast, vacuolar glucose accumulated stronger in western accessions while an increase of vacuolar fructose portions was indicative for cold acclimated eastern accessions. Vacuolar sucrose fractions together with relative amounts of glucose in plastids and cytosol were correlated with plants at 22°C.

**Fig. 4:**
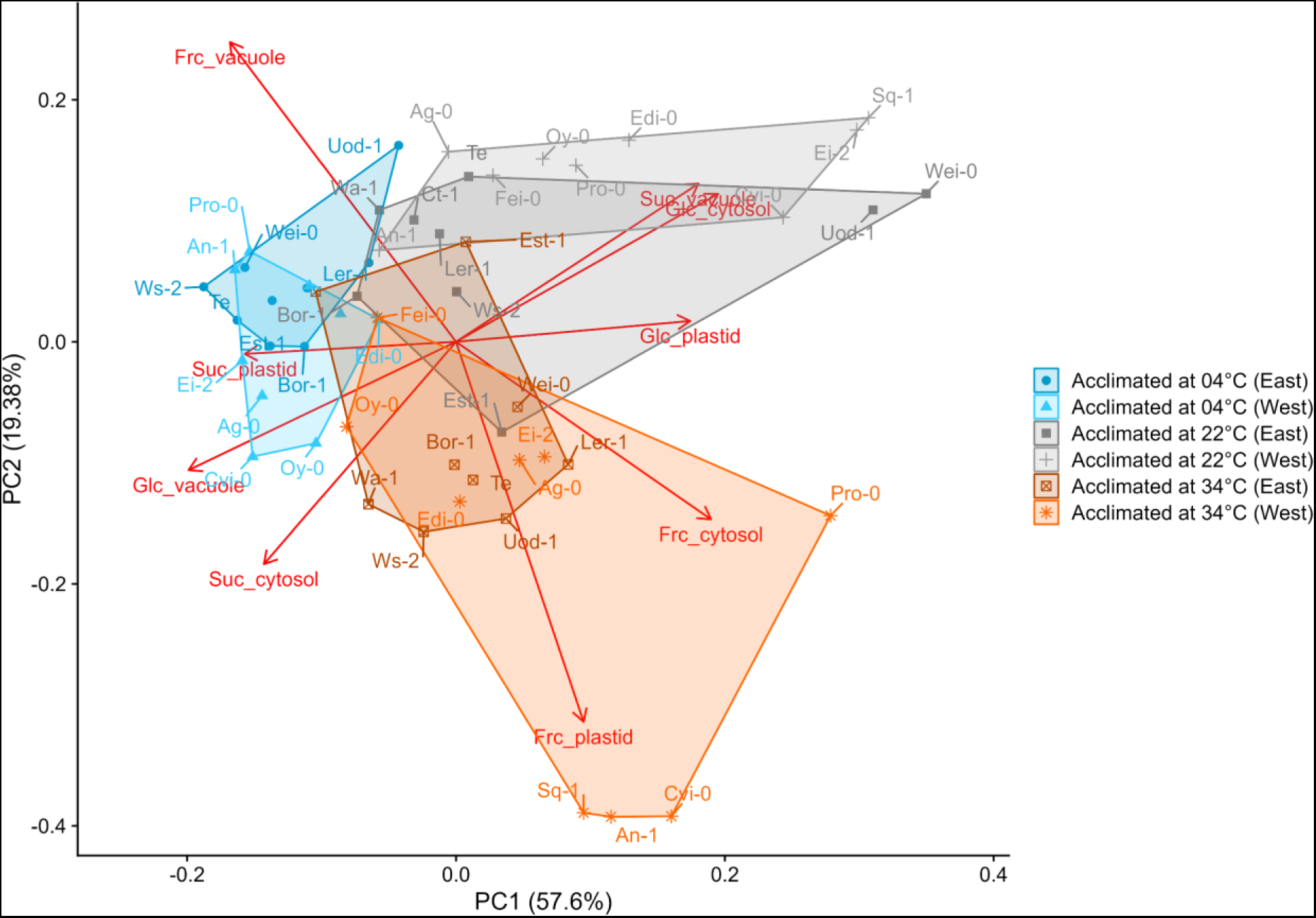
Natural variation of subcellular carbohydrate compartmentation during heat and cold acclimation. Dots represent scores of accessions acclimated at different temperatures as indicated by the color (grey: 22°C, blue: 4°C, orange: 34°C). Loadings represent relative carbohydrate distribution across plastids, cytosol and vacuole (n = 3). Classification of eastern and western accessions refers to scaled ETR residuals of heat acclimated plants (see Figure 1).

To estimate the impact of physiologically relevant subcellular metabolite concentrations on stability of photosynthesis, effective subcellular concentrations were calculated from relative metabolite distributions, their absolute amounts and assumptions about compartment volumes (Nägele and Heyer, 2013). The volume of plastids was estimated as 20 % of the total cellular volume. For cytosol, it was estimated to be 5 % and 75 % for the vacuole. The estimated compartment-specific concentration was then correlated to Fv/Fm, maximum ETR and residuals of scaled ETR (Fig. 5; Supplemental Tables S3 and S4).

**Fig. 5:**
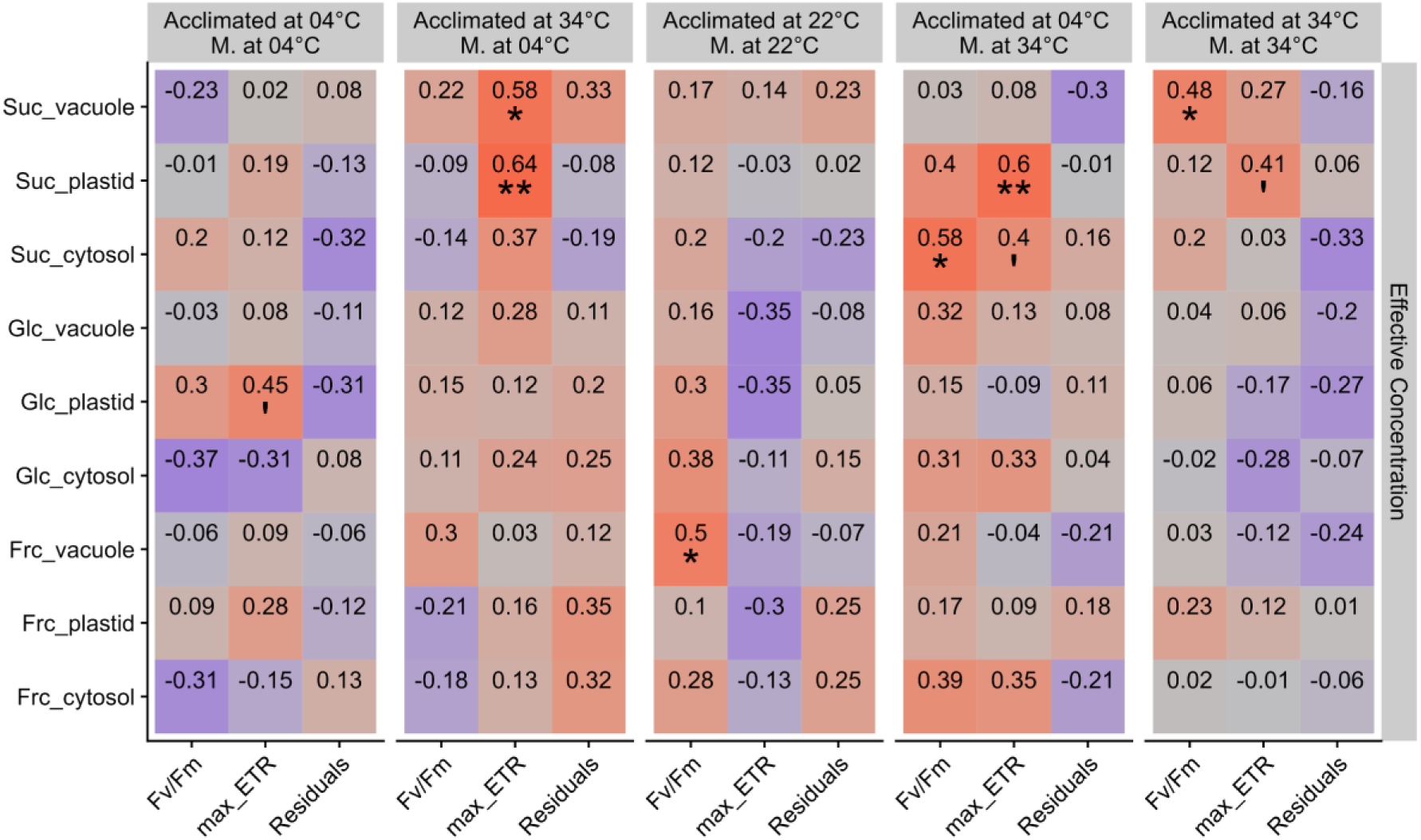
Correlation between photosynthetic efficiencies and subcellular metabolite concentrations. Numbers represent Spearman’s rank correlation coefficients. Columns relate to experiments to determine *Fv/Fm*, maximal ETR (*max_ETR*) and residuals of scaled ETR (*Residuals*), from left to right: Acclimated at 04°C, Measured at 04°C; Acclimated at 34°C, Measured at 04°C; Acclimated at 22°C, Measured at 22°C; Acclimated at 04°C, Measured at 34°C; Acclimated at 34°C, Measured at 34°C. Red color: positive correlation; purple color: negative correlation. Significance codes: ** 0.001 < p < 0.01; * 0.01 < p < 0.05; ‘ 0.05 < p < 0.1.

The strongest positive and most significant correlations were observed between the maximum ETR and the effective plastidial sucrose concentration for plants which were acclimated at 4°C and measured at 34°C, and *vice versa*. Also, vacuolar sucrose concentration correlated significantly with maximum ETR and Fv/Fm for heat acclimated plants, measured at 4°C and 34°C, respectively. A significant positive correlation was found for cytosolic sucrose concentration and Fv/Fm in plants acclimated to 4°C and measured at 34°C. In plants acclimated at 22°C, vacuolar fructose concentration was found to be significantly positive correlated to Fv/Fm.

To reveal how subcellular metabolite concentrations relate to the ETR residual-based classification of western and eastern accessions, Spearman’s rank correlation coefficients were also determined for both accession groups separately (Figure 6). Vacuolar sucrose concentrations were positively correlated with maximum ETR at 22°C in eastern accessions while, under fluctuating temperature (acclimated at 34°C è measured at 4°C and *vice versa*), plastidial sucrose concentrations significantly correlated with maximum ETR. This was also observed for cold acclimated western accessions. However, heat acclimated western accessions measured at 4°C showed a significant negative correlation of plastidial sucrose concentration with Fv/Fm which was contrasting eastern accessions. Also, plastidial fructose concentrations were negatively correlated with Fv/Fm in western accessions when heat acclimated plants were measured at 4°C. For heat acclimated western accessions, measured under 34°C, it was found that vacuolar sucrose concentration was negatively correlated with ETR residuals indicating a stabilizing function of photosynthesis. In general, those observations suggested that not only photosynthetic efficiencies in a fluctuating environment, but also subcellular metabolic acclimation strategies differed between accessions with distinct longitudinal origin.

**Fig. 6:**
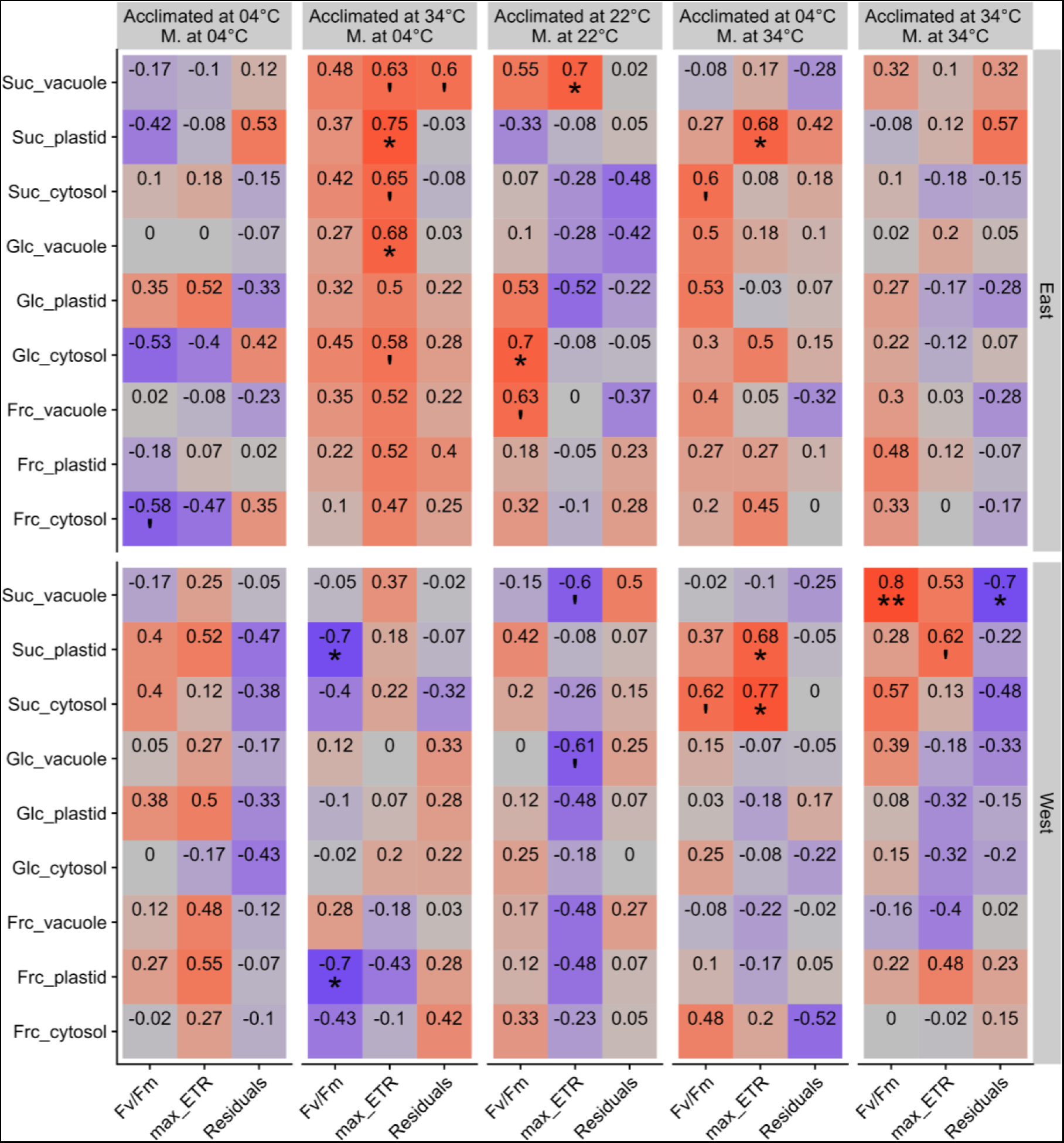
Correlation between photosynthetic efficiencies and subcellular metabolite concentrations of eastern (top panel) and western (lower panel) accessions. Numbers represent Spearman’s rank correlation coefficients. Columns relate to experiments to determine *Fv/Fm*, maximal ETR (*max_ETR*) and residuals of scaled ETR (*Residuals*), from left to right: Acclimated at 04°C, Measured at 04°C; Acclimated at 34°C, Measured at 04°C; Acclimated at 22°C, Measured at 22°C; Acclimated at 04°C, Measured at 34°C; Acclimated at 34°C, Measured at 34°C. Red color: positive correlation; purple color: negative correlation. Significance codes: ** 0.001 < p < 0.01; * 0.01 < p < 0.05; ‘ 0.05 < p < 0.1.

## 4. Discussion

Changing temperature regimes belong to typical environmental dynamics which plants are exposed to on different time scales. Already within a diurnal cycle, temperature changes might comprise large amplitudes. Comparing temperature amplitudes over recent decades has revealed an upward trend of minimum and maximum temperatures and an increased probability of extreme events in the temperate climate zone (Ouyang et al., 2023). Plant physiological and molecular consequences of such a dynamic environment can be estimated by analyzing temperature stress response and acclimation output. Immediate adjustment of photosynthesis and metabolism to environmental stressors, e.g., significant temperature fluctuation, represents a prerequisite for plant survival and long-term acclimation responses (Kosova et al., 2011). While forward genetic screens, QTL mappings and reverse genetic approaches have essentially contributed to the understanding of plant-environment interactions (Chinnusamy et al., 2003; Panter et al., 2019), the study of natural variation of traits has further contributed important insights. For example, comparing freezing tolerant to freezing sensitive accessions revealed a central role of the C-repeat binding factors (CBFs) in adjusting a low-temperature metabolome (Cook et al., 2004). The study of transcriptomes and metabolomes of natural accessions has further revealed negative regulators of freezing tolerance, e.g., auxin-and cytokinin-induced response regulator genes (Hannah et al., 2006). Simulating naturally occurring low temperature profiles during early vegetative growth showed that seedling growth differs significantly between *Arabidopsis* accessions originating from northern and southern latitudes which was suggested to reflect local adaptation mechanisms (Clauw et al., 2022). This was supported by substantial reprogramming of the transcriptome and primary metabolome resulting in a differential metabolome plasticity which was found to be negatively correlated to maximum temperature of the first quarter of the year at the respective natural habitat (Weiszmann et al., 2023). Also, in the present study, climate data of the first quarter of the year over almost seven decades were found to correlate negatively with fluctuations of photosynthetic (linear) ETR (see Fig. 1). Here, measurements at elevated temperature were considered because they showed least ETR residuals under temperature fluctuations which suggests the highest photosynthetic acclimation output across all experiments. Hence, the observed correlation output with climate data suggests that plants which originate from habitats with high temperature fluctuation during the first quarter of the year show more efficient photosynthetic acclimation than accessions from habitats with low temperature fluctuation. This fell together with a longitudinal gradient of the chosen accessions. However, maximum quantum yield of PSII was found to be stable across the full longitudinal range which indicates that these conditions are not selective enough to neither prevent growth of eastern accessions in western habitats nor *vice versa*.

The ratio of the insoluble storage compound starch to soluble carbohydrates has been found earlier to indicate differential metabolic reprogramming due to temperature changes across natural accessions (Klotke et al., 2004; Guy et al., 2008; Nagler et al., 2015). Here, starch was observed to accumulate stronger in cold acclimated western accessions while soluble carbohydrates were associated to eastern accessions. Although neither cold, freezing nor heat tolerance levels were quantified in the present study, due to differential ETR residuals under heat it might be speculated that western accessions tendentially represent more heat tolerant accessions while eastern accessions might be cold adapted. If this assumption pertained, reduced starch accumulation in eastern accessions might be due to increased starch degradation capacities to support accumulation of soluble carbohydrates and other cryoprotective compounds (Kaplan et al., 2006; Guy et al., 2008; Sicher, 2011). During heat exposure, starch amounts were found to decrease in most accessions which is in line with previous findings, and a reasonable explanation might be the inhibition of ADP-glucose pyrophosphorylase and starch synthesis than increased capacities of starch degradation (Geigenberger et al., 1998; Awasthi et al., 2014). Interestingly, some accessions showed less heat sensitive starch metabolism (e.g., Fei-0, Oy-0 or Sq-1) and it might be interesting to analyze differences in starch degradation pathways among these accessions in future studies.

As a result of increased starch degradation and sucrose accumulation under heat, many accessions displayed an increased sucrose-to-starch ratio (for an overview, see Fig. 7). This effect was at least as strong as for cold acclimated plants across all accessions and might be explained by decreased invertase expression (Prasch and Sonnewald, 2013). Subcellular analysis additionally revealed that, at low temperature, a significant fraction of sucrose was accumulating in the plastids which would preserve it from being cleaved by cytosolic and/or vacuolar invertases (Klotke et al., 2004; Weiszmann et al., 2018). While invertases have also been shown to occur in plastids (Vargas et al., 2008), their activity might be much lower than in other compartments. We have shown earlier that plastidial accumulation of sucrose represents a conserved cold acclimation strategy in *Arabidopsis* (Nägele and Heyer, 2013), maybe to support stabilization of thylakoid membranes and photosynthesis similar as raffinose (Knaupp et al., 2011) or to balance carbon allocation throughout the cell (Patzke et al., 2019). As indicated by the cold-induced accumulation of hexoses and depletion of sucrose in the vacuole, vacuolar sucrose cleavage, catalyzed by invertases, might represent another stabilization mechanism of photosynthesis as previously suggested (Weiszmann et al., 2018). Remarkably, this pattern was observed for all accessions, but eastern accessions accumulated fructose while western accessions rather accumulated glucose in the vacuole (Fig. 7). This finding indicates that not only sucrose biosynthesis and cleavage but also the subsequent metabolism of cleavage products glucose and fructose shows significant natural variation. This might be due to differential regulation of fructokinase and hexokinase enzymes. Hexokinase 1 (AtHXK1) catalyzes the phosphorylation of glucose and has recently been shown to improve drought and heat tolerance when overexpressed in potato (Lehretz et al., 2021). Further, AtHXK1 is a central conserved sugar sensor which might directly affect photosynthesis and transpiration by controlling stomatal aperture (Moore et al., 2003; Granot and Kelly, 2019). Fructokinase catalyzes the carbon flow into glycolytic pathway and the tricarboxylic acid cycle to provide substrates for amino acid biosynthesis and mitochondrial respiration (Pego and Smeekens, 2000). Hence, natural variation of cold and heat acclimation capacities may, among others, be due to differential hexose phosphorylation capacities which would also have impact on respiratory acclimation to a changing temperature regime (Talts et al., 2004; Atanasov et al., 2020).

**Fig. 7:**
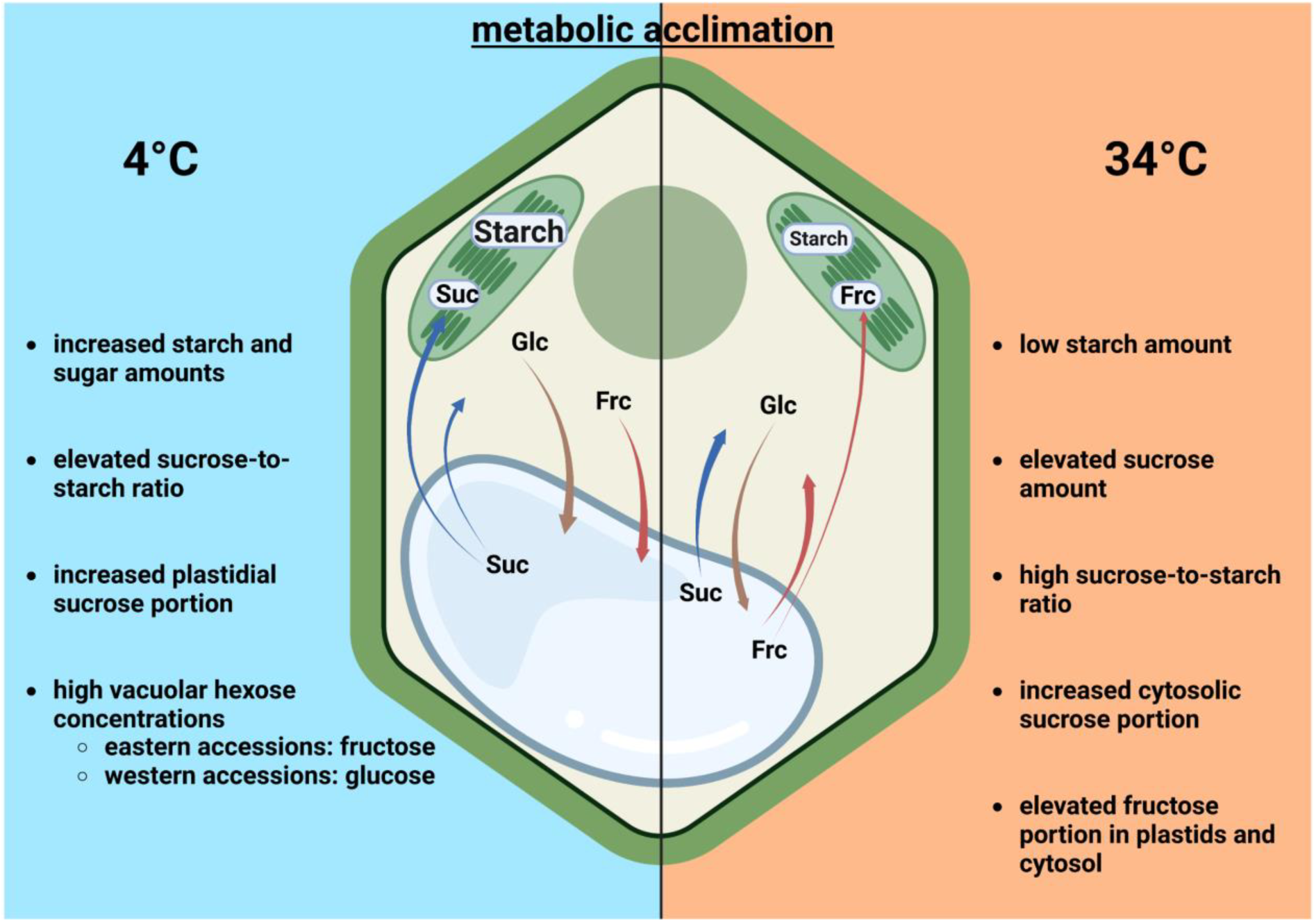
Comparing natural variation of metabolic acclimation to low and elevated temperature. Comparisons relate to non-acclimated plants (22°C). Arrow sizes indicate effect strength, arrow color refers to different sugars. Suc – sucrose (blue arrows); Glc – glucose (brown arrows); Frc – fructose (red arrows). Created with BioRender.com.

An increasing body of evidence has revealed that, to comprehensively understand regulation of photosynthesis and metabolic processes involved in plant temperature response, subcellular data needs to be considered (Hoermiller et al., 2017; Hurry, 2017; Höhner et al., 2021). Yet, estimating effective *in vivo* metabolite concentrations remains challenging not only because of laborious sample fractionation but also due to lacking information about organellar volumes and their temperature-dependent dynamics. Particularly, for quantitative analysis of compartment-specific reaction rates and metabolic fluxes such information becomes essential because compartmentalization of metabolism directly affects enzymatic activity and function, thus altering regulation and fluxes within a metabolic network (Szecowka et al., 2013; Herrmann et al., 2021; Nägele, 2022). Data of the present study provides further evidence for the importance of estimating subcellular effective concentrations which enabled the interpretation of physiological effects on photosynthesis. Comparing correlations between subcellular sugar concentrations and photosynthetic efficiencies indicated the importance of splitting the accessions into east and west. Even under ambient temperature, such a split was essential to reveal positive and negative correlation of vacuolar sucrose concentrations with Fv/Fm and maximum ETR in eastern and western accessions, respectively. Hence, uncovering such effects supports our hypothesis of a physiologically relevant classification due to geographical origin and habitat information.

Finally, many significant correlations between subcellular sugar concentrations and photosynthetic efficiencies were only observed under fluctuating temperature which suggests that a subcellular metabolic adjustment during temperature acclimation results in a metabolic state which stabilizes photosynthesis against further environmental deflections. This hints towards the necessity of rethinking the definition of plant (temperature) acclimation which might rather consider the plasticity of metabolism towards environmental changes than its absolute configuration.

## Author contributions

J.H. performed experiments and data evaluation, developed the R app (NAFalyzer) and wrote the paper. D.D. performed experiments and supported data evaluation. L.S., C.S. and A.K. performed experiments. V.B. developed the R app (NAFalyzer). T.N. conceived the study, supported data evaluation and wrote the paper.

## Acknowledgments

We would like to thank all members of Plant Evolutionary Cell Biology at the Faculty of Biology, LMU Munich, for many fruitful discussions. We thank Andreas Klingl from Plant Development, LMU Munich, for support of CS. We would also like to thank Svenja Eberlein and Russell Castelino for assisting in PAM measurements.

## Supplemental Information

**Fig. S1. Natural variation of maximum quantum yield of PSII and linear electron transport rate. (A)** Maximum quantum yield of PSII (Fv/Fm) for individual accessions after temperature acclimation. Fv/Fm was determined at different temperatures (measured at 4°C, 22°C or 34°C). **(B)** Maximum quantum yield of PSII (Fv/Fm) for eastern (orange) and western (blue) accessions after temperature acclimation. Fv/Fm was determined at different temperatures (measured at 4°C, 22°C or 34°C). **(C)** Linear ETR for individual accessions as a function of PAR. ETR was determined at different temperatures (measured at 4°C, 22°C or 34°C). **(D)** Linear ETR for for eastern (orange) and western (blue) accessions as a function of PAR. ETR was determined at different temperatures (measured at 4°C, 22°C or 34°C). Parameters were determined after growth for 5 weeks under ambient growth conditions before and after being exposed to a temperature treatment of either 4, 22, or 34°C for 4 days (lines represent means; n ≥ 5). A list of all accession information is provided in the supplements (Supplementary Table S1).

**Fig. S2. Correlation of Fv/Fm and ETR residuals.** Maximum quantum yield of PSII (Fv/Fm; ordinates) was correlated with ETR residuals (abscissae) for each accession (color coded). Kendall’s rank correlation coefficient (cor) and significance (p-value) are shown for each correlation analysis. Columns described acclimation temperatures (4°C, 22°C, 34°C; left-to-right), rows indicate measurement temperature (4°C, 22°C, 34°C; top-down)

**Fig. S3. Metabolite amounts of natural accessions.** (**A**) Starch amount in C6 equivalents, (**B**) Sucrose amount in C6 equivalents, (**C**) Glucose amount, and (**D**) Fructose amount, all normalized to dry weight (gDW-1). (**E**) Sucrose-to-starch ratio, calculated with C6 equivalents. Plants were grown for 5 weeks before being transferred to either 4°C (blue bars), 22°C (gray bars) or 34°C (orange bars) for 4 days. n ≥ 5.

**Table S1. Information about natural accessions.** The table contains information about latitude and longitude of the geographical origin. Maximum temperature (max_temp), minimum temperature (min_temp), mean temperature (mean_temp), deviation of recorded mean temperature (d_temp) and mean precipitation (mean_precip) are provided for the first quarter of the year (1950 – 2019; https://climatecharts.net/).

**Table S2. Pearson correlation coefficients of subcellular sugar distribution, climate and geographical data of natural accessions.** Climate data was used from the first quarter of the year, i.e., January to March. *M temperature*: mean temperature; *Δ temperature*: difference between the highest and lowest average temperature of the individual months. Significance codes: ** 0.001 < p < 0.01; * 0.01 < p < 0.05; ‘ 0.05 < p < 0.1.

**Table S3. Experimentally determined electron transport rates, maximum quantum efficiencies of PSII and residuals.**

**Table S4. Metabolite amounts and relative subcellular distributions.**

