## Supplemental Table S2 for "Temperature acclimation of photosynthesis and carbohydrate metabolism are related to the geographical origin of *Arabidopsis thaliana*"

**Table S2. Pearson correlation coefficients of subcellular sugar distribution, climate and geographical data of natural accessions.** Climate data was used from the first quarter of the year, i.e., January to March. *M temperature*: mean temperature; *Δ temperature*: difference between the highest and lowest average temperature of the individual months. Significance codes: ** 0.001 < p < 0.01; * 0.01 < p < 0.05; ‘ 0.05 < p < 0.1.

| | Condition | *M* temperature | Δ temperature | Longitude | Latitude |
|---|---|---|---|---|---|
| Fructose plastid | 4°C | 0.61 ** | -0.39 | -0.55 * | -0.6 ** |
| Fructose cytosol | 4°C | -0.2 | -0.17 | -0.07 | 0.31 |
| Fructose vacuole | 4°C | -0.1 | 0.34 | 0.32 | -0.01 |
| Fructose plastid | 22°C | -0.28 | 0.28 | 0.33 | 0.12 |
| Fructose cytosol | 22°C | 0.2 | -0.1 | -0.36 | -0.24 |
| Fructose vacuole | 22°C | -0.1 | 0.01 | 0.23 | 0.18 |
| Fructose plastid | 34°C | 0.38 | -0.34 | -0.46 ‘ | -0.34 |
| Fructose cytosol | 34°C | 0.25 | -0.13 | -0.43 ‘ | -0.3 |
| Fructose vacuole | 34°C | -0.36 | 0.27 | 0.51 * | 0.37 |
| Glucose plastid | 4°C | 0.21 | -0.22 | 0.01 | -0.13 |
| Glucose cytosol | 4°C | -0.31 | 0.2 | 0.11 | 0.31 |
| Glucose vacuole | 4°C | 0.22 | -0.11 | -0.1 | -0.24 |
| Glucose plastid | 22°C | 0.23 | -0.04 | -0.29 | -0.26 |
| Glucose cytosol | 22°C | 0.3 | -0.13 | -0.43 ‘ | -0.29 |
| Glucose vacuole | 22°C | -0.29 | 0.12 | 0.41 ‘ | 0.29 |
| Glucose plastid | 34°C | 0.03 | -0.1 | -0.26 | -0.07 |
| Glucose cytosol | 34°C | 0.2 | -0.23 | -0.42 ‘ | -0.25 |
| Glucose vacuole | 34°C | -0.14 | 0.19 | 0.38 | 0.19 |
| Sucrose plastid | 4°C | -0.17 | 0.38 | 0.28 | 0.02 |
| Sucrose cytosol | 4°C | 0.07 | -0.47 ‘ | -0.11 | 0 |
| Sucrose vacuole | 4°C | 0.04 | 0.15 | -0.07 | -0.01 |
| Sucrose plastid | 22°C | 0.03 | 0.23 | 0.1 | -0.2 |
| Sucrose cytosol | 22°C | -0.06 | 0.16 | 0.36 | 0 |
| Sucrose vacuole | 22°C | 0.05 | -0.2 | -0.33 | 0.05 |
| Sucrose plastid | 34°C | 0.1 | -0.05 | -0.08 | -0.04 |
| Sucrose cytosol | 34°C | -0.17 | 0.09 | 0.18 | 0.23 |
| Sucrose vacuole | 34°C | 0.14 | -0.07 | -0.15 | -0.21 |
