## Supplementary figures and images for "Temperature acclimation of photosynthesis and carbohydrate metabolism are related to the geographical origin of *Arabidopsis thaliana*"

### Supplemental Figure S1

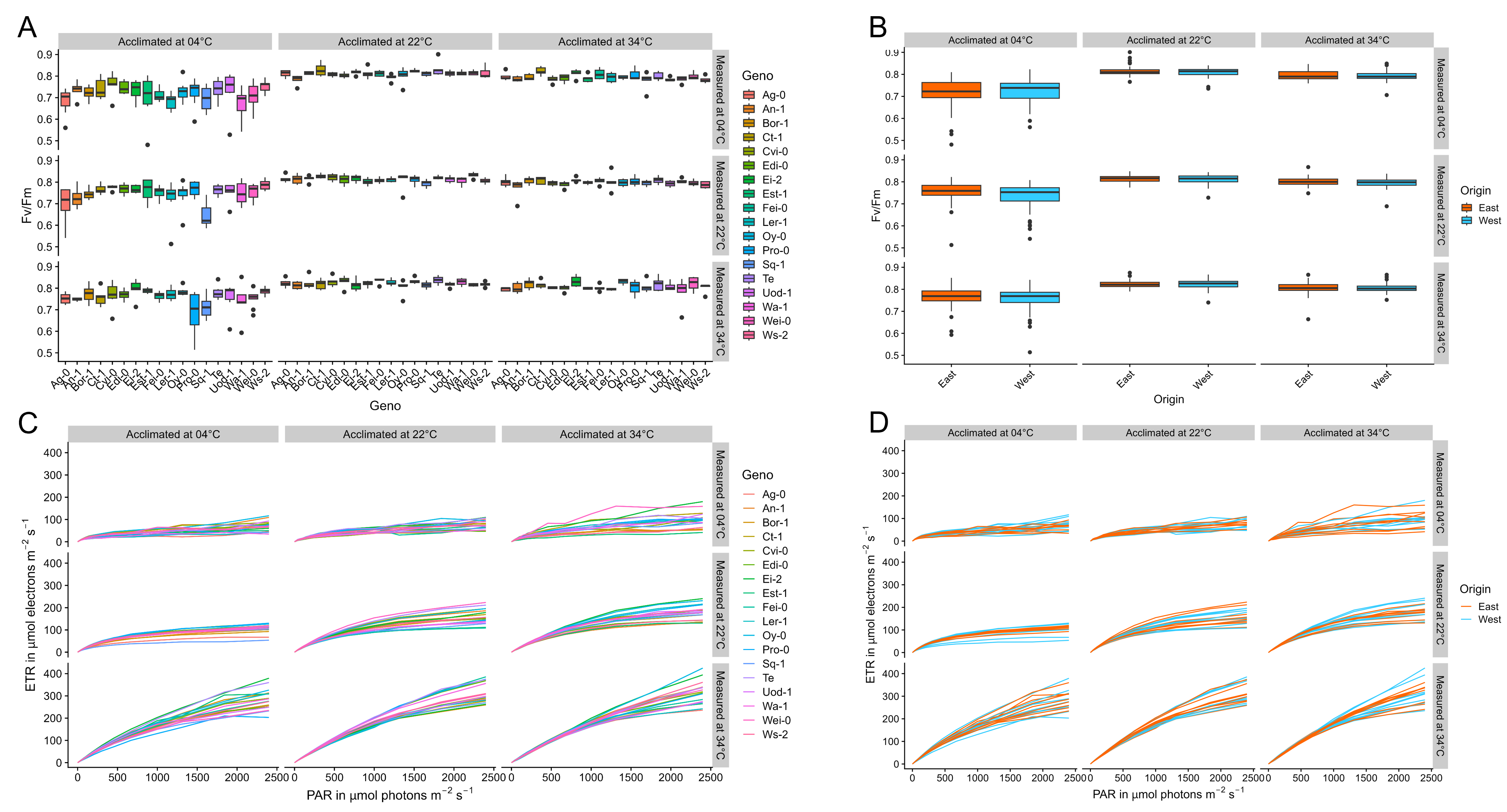

### Supplemental Figure S2

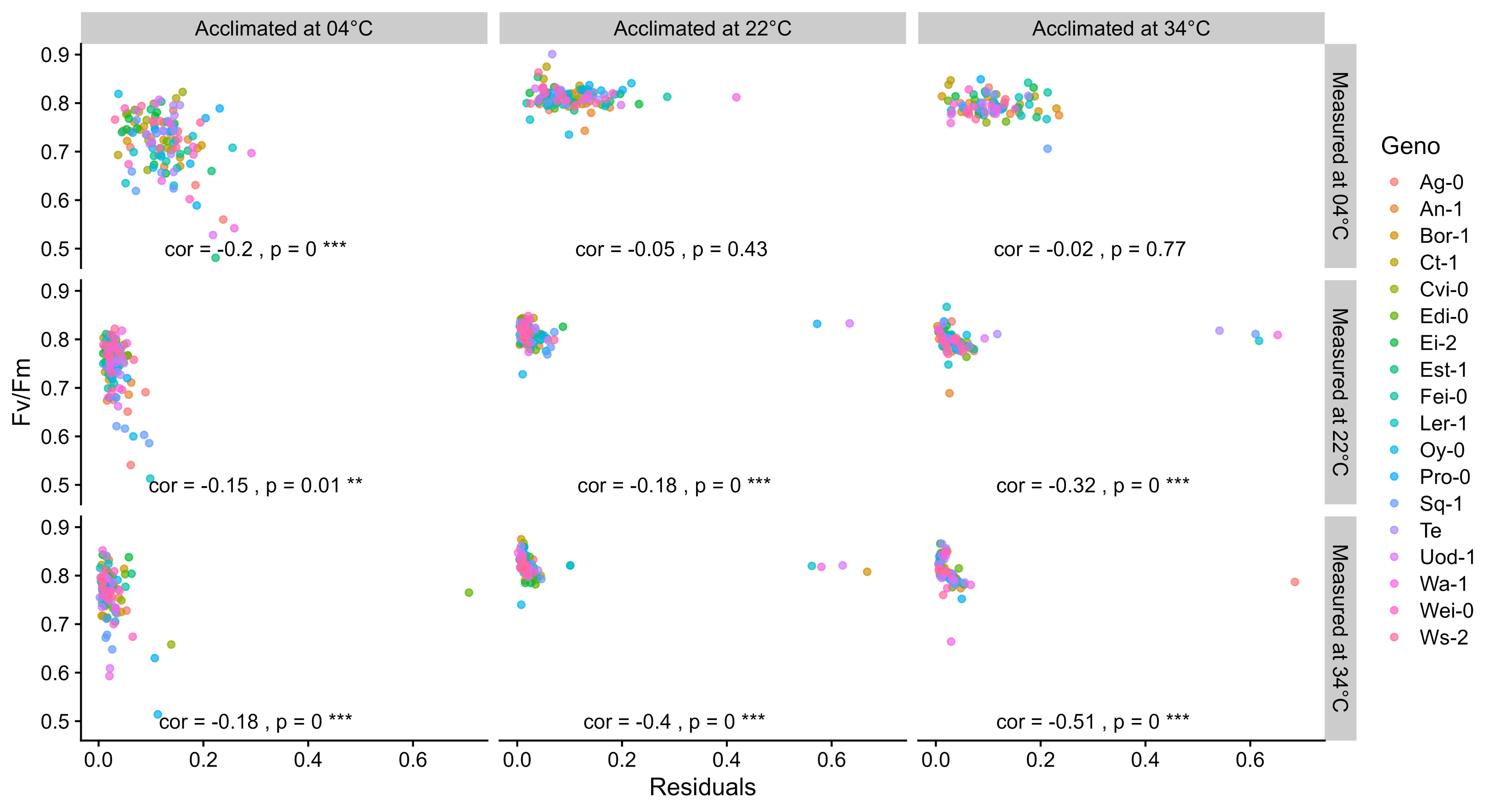

### Supplemental Figure S3

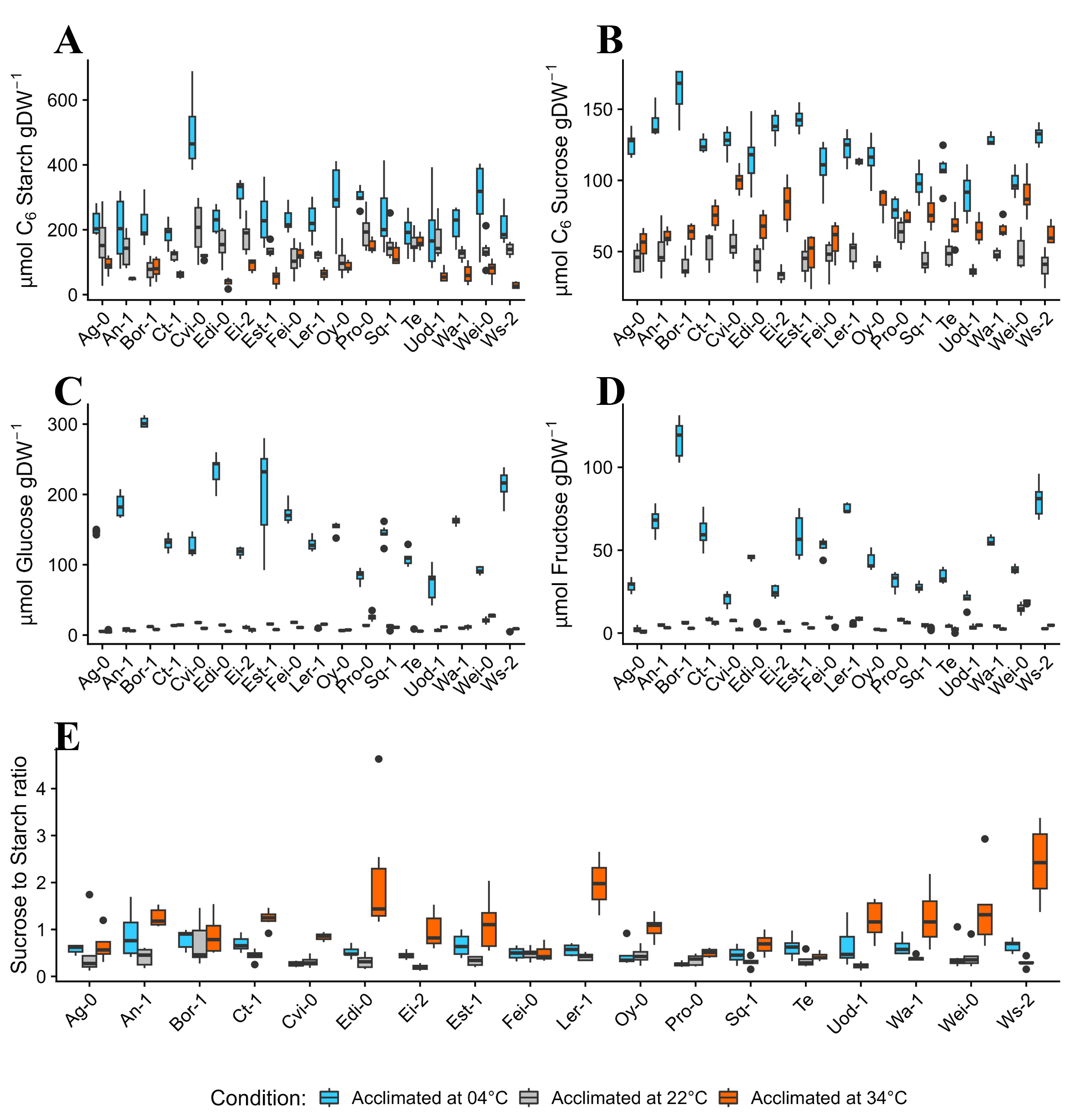
